# Prior hypotheses or regularization allow inference of diversification histories from extant timetrees

**DOI:** 10.1101/2020.07.03.185074

**Authors:** Hélène Morlon, Florian Hartig, Stéphane Robin

## Abstract

Phylogenies of extant species are widely used to study past diversification dynamics^1^. The most common approach is to formulate a set of candidate models representing evolutionary hypotheses for how and why speciation and extinction rates in a clade changed over time, and compare those models through their probability to have generated the corresponding empirical tree. Recently, Louca & Pennell^2^ reported the existence of an infinite number of ‘congruent’ models with potentially markedly different diversification dynamics, but equal likelihood, for any empirical tree (see also Lambert & Stadler^3^). Here we explore the implications of these results, and conclude that they neither undermine the hypothesis-driven model selection procedure widely used in the field nor show that speciation and extinction dynamics cannot be investigated from extant timetrees using a data-driven procedure.

## Main text

Louca & Pennell^2^ consider the homogeneous (i.e. lineage-independent) stochastic birth-death process of cladogenesis traditionally used in macroevolution to test hypotheses about how and why rates of speciation (birth, *λ*) and extinction (death, *μ*) have changed over time *t*. They show that for any given time-dependent speciation function *λ* > 0 and extinction function *μ* ≥ 0, there exists an infinite set of alternative functions *λ*^*^ > 0 and *μ*^*^ ≥ 0 such that the probability distribution of extant trees under the corresponding birth-death processes M and M^*^ is identical. Consequently, M or M^*^ yield identical likelihood values for any given empirical tree. This identifiability issue is certainly both interesting and unfortunate, but what are its implications for phylogenetic-based diversification analyses?

We consider this question against two alternative philosophies to conducting science. The first is a hypothesis-driven research approach, in which a small number of alternative ideas about the underlying mechanism are compared against data^4^. An example is the foundational study of Nee *et al*. (1992)^5^, who examined one of the first molecular phylogenies of birds, and demonstrated that it was incompatible with a constant-rate diversification model. Grounded in Simpson’s evolutionary theory of adaptive radiations^6^, they then hypothesized that rates of cladogenesis in birds might be affected by niche-filling processes. Finding that a diversity-dependent model indeed fitted their data better, they concluded that diversity-dependent cladogenesis was a more plausible scenario to explain the diversification of birds.

This hypothesis-driven approach has inspired more than 30 years of research in phylogenetic diversification analyses^1^. Exponential time-dependencies have been used, for example, to mimic early burst patterns expected from adaptive radiation theory^6^, or as an approximation to diversity-dependent cladogenesis^7^. In the context of the environment-dependent models mentioned by Louca & Pennell^2^, functional hypotheses have often been derived from foundational theories of biodiversity, such as the metabolic theory of biodiversity^8^ and MacArthur & Wilson’s theory of island biogeography^9^. Phenomenological models, such as simple linear time- or environmental-dependencies, have indeed also been used, but typically either as null models^7^ or as the simplest way to model the effect of an explanatory environmental variable on evolutionary rates^8,10^. The primary goal of this research, however, has been to fit, test and compare diversification scenarios that were defined *a priori* to reflect different evolutionary hypotheses.

Louca & Pennell’s congruent models M^*^, on the other hand, do not correspond *a priori* to any evolutionary hypotheses, and would never be considered in a well-conducted hypothesis model selection procedure in the first place^4^. Fortunately, as Louca & Pennell^2^ admit, most diversification hypotheses that are compared by evolutionary biologists are distinguishable from extant trees, and the ongoing effort to integrate fossil information provides even brighter perspectives^11^. The existence of a large number of congruent models therefore poses no direct challenge to the traditional hypothesis-driven research approach. The only possible concern is the question of model selection consistency: if the true model is not in the set of considered models, do we select the correct hypothesis? This question has not been answered one way or the other and would require thorough investigation in future research.

By considering all possible diversification functions, Louca & Pennell^2^ implicitly subscribe to a different method of scientific discovery, where the goal is to learn *λ* and *μ* from the data alone. After finding that these quantities are not simultaneously identifiable, even for infinitely large phylogenies, they suggest to instead estimate identifiable quantities such as the pulled speciation rate *λ_p_* or pulled diversification rate *r_p_*. There are certainly advantages of the pulled rates, but estimating them from limited-size phylogenies is still a challenging task (SI S. 1). One could address the problem by regularization, but then the question arises why we should not directly use regularization to handle the identifiability issue around *λ* and *μ*. Indeed, a wide body of statistical regularization techniques exist to deal with unidentifiability issues^12^, such as shrinkage^13^, smoothing^14^ or the use of Bayesian priors. Louca & Pennell^2^ do not address the possibility to use such statistical regularization methods, although these methods have already been used successfully for inference of diversification rates^15,16^ (SI S.2).

To provide a concrete example of these points, we perform an analysis of the diversification of the Madagascan vangas in the logic that would be applied in the field^17^, but simplified for illustrative purposes. We hypothesize that diversification followed an ‘Early Burst’ pattern^18^, with fast speciation at the origin of the group and subsequent slowdown, rather than constant-rate diversification. The Early Burst pattern, related to the idea of adaptive radiations^6^, is modeled by an exponential decay of the speciation rates through time, used as an approximation of diversity-dependence. We also consider the hypothesis that a substantial number of extinction events occurred during the diversification of this group. Among the four corresponding models, the model with exponential change and no extinction (M) is best supported by the data (see Table 1 & Supplementary Information). M is characterized by *λ*(*t*) = *λ*_0_*e^αt^* and *μ*(*t*) = 0 where *t* is measured from the present to the past, *λ*_0_ = 0.018 is the estimated present-day speciation rate, and *α* = 0.1 measures the estimated speed of time change. A positive *α* reflects a decline in speciation rate from the origin of the group to the present (Fig. 1). We conclude that the hypothesis of Early Burst diversification with negligible extinctions is the most likely of the four hypotheses we considered.

**Figure 1.**
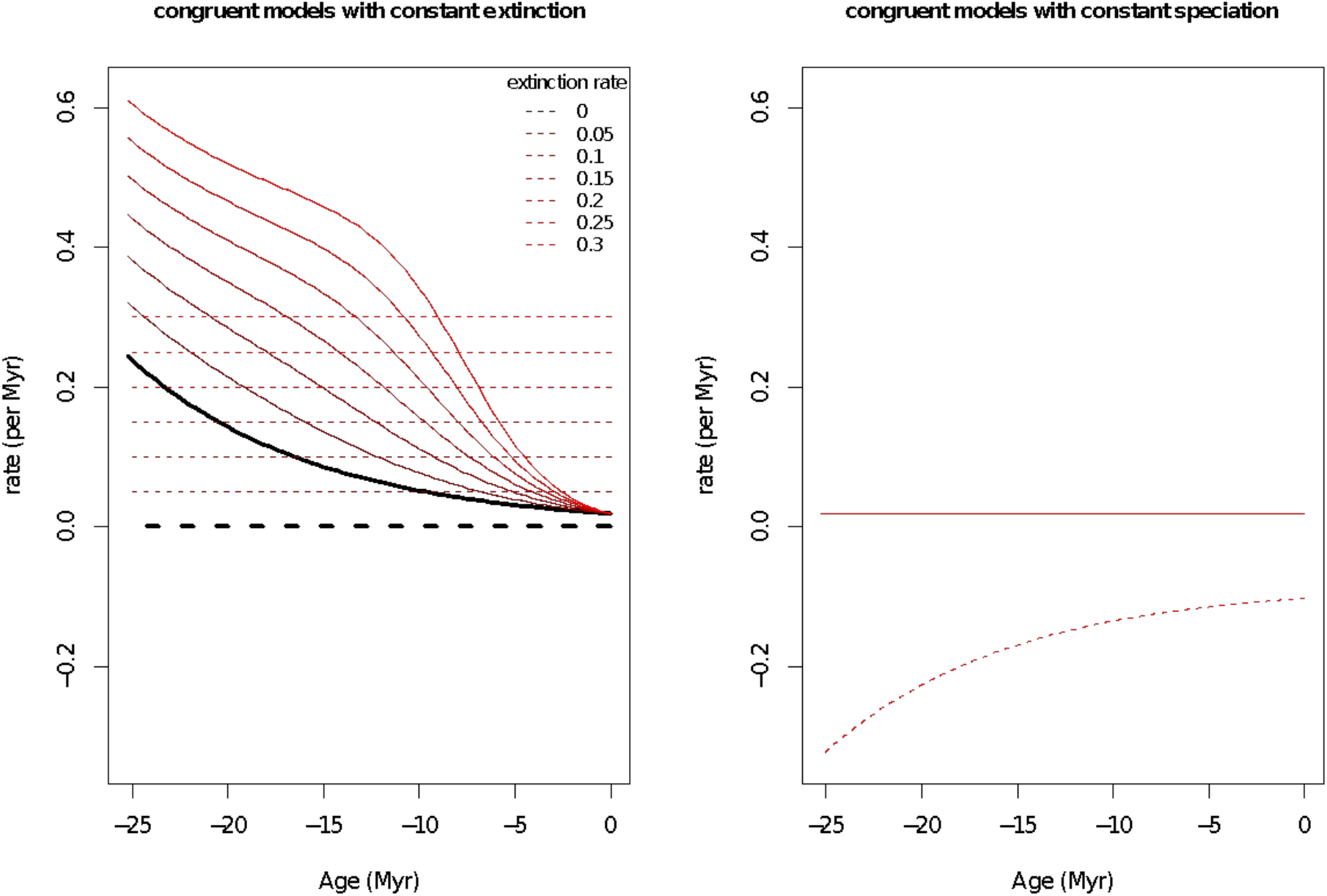
Diversification of the Madagascan vangas as inferred from congruent models. The black curves on the left panel represent the dynamics of speciation (plain line) and extinction (dashed line) corresponding to our best-fit model M (exponential decline in speciation rate, non-significant extinctions). The colored curves illustrate the rate dynamics of congruent models obtained by: fixing increasing values of a constant extinction rate (left panel, M_1_^*^) and fixing the speciation rate to *λ*_0_ (right panel, M_2_^*^). Note that M1* infers a speciation rate decline regardless of the assumed extinction rate.

**Table 1.** Diversification models fitted to the phylogeny of the Madagascan vangas.

| Model | nb | Log L | AICc | $\Delta AIC$ | $\lambda_0$ | $\alpha$ | $\mu_0$ |
| --- | --- | --- | --- | --- | --- | --- | --- |
| Exponential change of speciation rate, no extinction (M) | 2 | -71.29 | 147.21 | 0 | 0.018 | 0.102 | - |
| Exponential change of speciation rate, constant extinction | 3 | -71.04 | 149.4 | 2.19 | 0.025 | 0.117 | 0.077 |
| Constant speciation, no extinction | 1 | -76.09 | 154.37 | 7.17 | 0.06 | - | - |
| Constant speciation, constant extinction | 2 | -76.09 | 156.80 | 9.6 | 0.06 | - | 3.39e-09 |

nb denotes the number of parameters. LogL stands for the maximum log-likelihood, AICc for the second order Akaike Information Criterion<sup>4</sup>, and $\Delta AIC$ for the difference in AICc between the model and the best model in the set. Models are ranked based on their AICc support. $\lambda_0$ is the estimated speciation rate at present, $\alpha$ is the estimated rate of change of speciation with time (a positive $\alpha$ reflects a rate decline), and $\mu_0$ is the estimated extinction rate. Fits were performed using the *fit\_bd* function from RPANDA<sup>21</sup> using the “crown” conditioning and a sampling fraction of 1 (the tree of the Madagascan vangas is complete<sup>17</sup>).

We now follow Louca & Pennell^2^ by solving Eq. 2 for models congruent to our best model M. First, we choose the extinction function to be a constant *μ*_1_ ^*^(*t*) = *μ*_0_ and compute *λ*_1_^*^(*t*). Second, we choose the speciation function to be a constant *λ*_2_^*^(*t*) = *λ*_0_ and compute *μ*_2_^*^(*t*). We find (SI; see the corresponding inferred dynamics in Fig. 1):

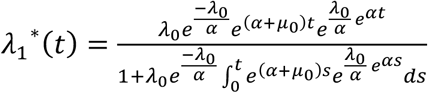

and

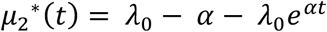

The biological interpretation of these additional models and of their parameters is not obvious. The equation for *μ*_2_^*^ looks more interpretable at first, but it expresses the temporal change and the extinction rate at present through the same parameter *α*, which means that a positive extinction rate at present (*α* < 0) will force extinction rates to decline over time. Here M_2_^*^ infers negative extinction rates, and is therefore not plausible (Fig. 1). M_1_^*^ infers a decline in speciation rate from the origin of the group to the present for extinction rates *μ*_0_ ranging from at least 0.05 to 0.3, consistent with our previous results (Fig. 1). While rate estimates do vary substantially, the general temporal trend is preserved.

Finally, we fit *r_p_* directly with spline functions, as suggested by Louca & Pennell^2^ (SI S.3), and deduce *λ* (or *μ*) when assuming *μ* (or *λ*) constant. General diversification trends are consistent with those found above; finer dynamics depend on the choices made to estimate *r_p_* and suggest interesting new hypotheses to test (SI S.3). Here we chose *λ* or *μ* to be constant to simplify the problem. Data-driven approaches have relaxed this hypothesis while still estimating global temporal tendencies accurately, based on the reasonable prior belief that rates don’t change much in a small amount of time^15^. We expect such or similar regularization approaches to eliminate the pathological cases with markedly different diversification histories shown in Louca & Pennell^2^.

There are useful results in Louca & Pennell’s paper. The pulled diversification rate is difficult to interpret biologically, yet realizing that models are only distinguishable by this quantity is important, and the approach might provide an interesting way to generate new hypotheses. The implications of these results for diversification analyses, however, are largely overinterpreted, mainly because the constraints imposed by the practice of hypothesis-driven research, prior knowledge, and the possibilities to penalize complexity are not considered in the paper. A very similar identifiability issue occurred 10 years ago in population genetics, when it was shown that the widely-used allelic frequency spectrum (AFS) is consistent with a myriad of demographic histories^19^. Despite this insight, the AFS – along with other data sources – remains a prominent approach of demographic inference^20^. Identifiability issues naturally arise in approaches that try to infer the potentially unlimited complexity of historical processes from limited contemporary data, and this is why we work hypothesis-driven, develop regularization techniques, and integrate other data types. Louca & Pennell^2^ remind us that the hypotheses we formulate when we develop models influence the conclusions we draw, and that these conclusions should always be taken with a grain of salt; this is always a good reminder, but it does not compromise the current practice in the field.

## Acknowledgments

We thank Amaury Lambert and the Morlon (current and past) research team, in particular Ignacio Quintero, Isaac Overcast, Jonathan P. Drury, Odile Maliet, and Julien Clavel for discussions. We also thank Knud Jønsson for providing us with the Magagascan vanga tree. H. Morlon acknowledges funding from the ERC (grant ERC-CoG PANDA).

## Supplementary Information

### S.1 Fitting the pulled rates also requires model selection or regularization

In their S9, Louca & Pennell explain how to obtain non-parametric estimates of *λ_p_* and *r_p_* from empirical timetrees by maximum likelihood. We note that, as for any non-parametric approach, these estimates rely on two non-independent and somewhat arbitrary choices: a) the functional basis (e.g. piecewise-polynomial, wavelets, Fourier) and b) the form of the regularization (e.g. smoothness, sparsity). For phylogenies of a few hundred of species, which is the data size typical for the application of the homogenous rate models considered by Louca & Pennell, the final estimates will depend on both choices. In other words, although *λ_p_* and *r_p_* are theoretically (i.e. for infinitely large phylogenies) identifiable, in practice they must be estimated from limited data. Without fixed hypotheses or constraints on functional complexity, the exact estimation of the pulled rates can therefore still be challenging. For example, Louca & Pennell chose a spline between an arbitrary number of discrete times as the functional basis. As we illustrate below, the choice of the degree of the spline and of the number of discrete times can influence estimates (see S.3). Our point is that some constraints, either in the form of fixed hypotheses that are chosen *a priori*, or in the form of constraints on model complexity, are fundamentally unavoidable in the problem of inferring diversification histories from extant time trees.

### S.2 Regularization techniques can allow inference of diversification histories despite unidentifiability

Louca & Pennell write that “common model selection methods that are based on parsimony or ‘Occam’s razor’ (such as the Akaike information criterion) generally cannot resolve these issues” (i.e. issues of model congruence). We first note that the Akaike information criterion is designed to account for the number of parameters in a (parametric) model, while most data-driven approaches are non-parametric. In the latter case, other approaches that are regularly used in statistics and machine learning to deal with issues of unidentifiability and over-parametrization include the use of shrinkage^13^ or smoothing^14^ estimators, or of Bayesian priors. The authors don’t discuss these approaches, but we see no reason why they should not be considered in this case.

Regarding the use of model selection, two main arguments are provided by the authors for dismissing these techniques as a possible solution to the identifiability problem (section S2 in Louca & Pennell’s paper, the third argument is essentially the same as the first). The first argument is that “There is little reason to believe that the simplest scenario in a congruence class will be the one closest to the truth. Indeed, even if the true model is included in a congruence class, it will almost always be the case that there are both simpler and more complex scenarios within the same congruence class and, crucially, all of these alternative models remain equally likely even with infinitely large datasets.” Nothing about this statement is specific to birth-death models adjusted to timetrees, and thus we must conclude that the authors question the principle of parsimony (or Occam’s razor) in general. The authors are of course entitled to their opinion, as the principle of parsimony is not a mathematically provable law, but we note that this opinion contradicts centuries of thinking and experience from physics to machine learning, and from philosophy as well. If we follow the traditional thinking in science when presented with the situation highlighted by Louca & Pennell (i.e. scenarios with different degrees of complexity but equal likelihood), we should select the simplest scenario, because, although there is no guarantee that this scenario is the closest to the truth, the parsimony principle suggests that it is most probably true, all other things equal.

The second argument by Louca & Pennell is that the complexity of a diversification scenario cannot be quantified. Note that this argument is only relevant if one accepts the parsimony principle in the first place, or else complexity is irrelevant for model selection. We agree with Louca & Pennell that quantifying complexity (or its opposite: simplicity) is a hard question, and one that is not always well-addressed by simply counting the number of parameters (the number of parameters is in particular not meaningful in non-parametric data-driven approaches). However, this again is not specific to birth-death models adjusted to timetrees, and a wide body of statistical literature exists that deals with this problem^12^. A review of this literature is beyond the scope of this discussion, but there are various routes to quantifying or penalizing complexity, such as information-theoretical measures (in particular the principle of maximum entropy^22^), other Bayesian and frequentist model selection approaches, and shrinkage or regularization estimators that essentially amount to setting priors on the parameters or smoothness of a curve. While we agree that choosing among those methods has a degree of subjectivity, we see no reason why complexity cannot be quantified for diversification scenarios. It seems to us that the problem is very similar to the general situation in data-driven analysis (starting from linear regressions), where we can always build more complex models with equal likelihood, but know for various reasons (parsimony, bias-variance trade-off) that adding complexity brings disadvantages, and thus add penalizations or other regularization methods to balance the complexity of the explanation. There have already been a few studies that tested regularization approaches for inference of diversification rates^15,16,23^, but the study by Louca & Pennell highlights the need for more research in this field, to explore what regularization methods work best.

Louca & Pennell also allude to the fact that the problem highlighted in their study is independent of sample size. We agree that this makes the situation somewhat different from some other model selection and regularization problems, but we do not see how this is relevant in a practical data analysis, where the size of the timetree is fixed and typically not huge (in particular if we assume homogeneous rates), meaning that the problem at hand is to draw inference from limited data.

### S.3 Illustration with the Madagascan vangas

Here we begin by exploring models congruent to M, the model selected among the four models we considered in our illustrative hypothesis-driven model selection procedure on the Madagascan vangas tree. M is characterized by *λ*(*t*) = *λ*_0_*e^αt^* and *μ*(*t*) = 0 where *t* is measured from the present to the past, *λ*_0_ = 0.018 is the estimated present-day speciation rate, and *α* = 0.1 measures the estimated speed of time change.

We first choose the extinction function to be a constant *μ*_1_^*^(*t*) = *μ*_0_ and compute *λ*_1_^*^(*t*). Note that the corresponding model M_1_^*^ has 3 parameters (*λ*_0_, *α* and *μ*_0_) while M has only 2 (*λ*_0_ and *α*), such that it would not be selected over M based on Occam’s razor. M_1_^*^ would not be selected either if compared to the model ranked second in our analysis (exponential change of speciation rate and significant extinctions), as it has the same number of parameters and a lower likelihood (Table 1). We compute *λ*_1_^*^(*t*) using the solution to Eq.2 from Louca & Pennell^2^, provided in their SI (Eq. 39 & 40), with *η*_0_ = *λ*_0_ (here the sampling fraction *ρ* = 1, as the Madagascan vangas tree is complete^17^):

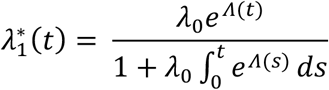

with

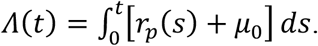

*r_p_*, the ‘pulled diversification rate’, is given by Eq. 1 from Louca & Pennell^2^:

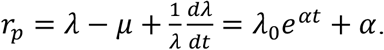

This gives:

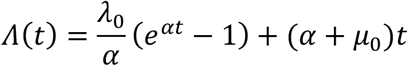

and

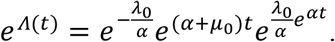

Finally:

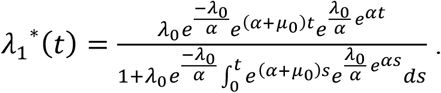

Second, we choose the speciation function to be a constant *λ*_2_^*^(*t*) = *λ*_0_ and compute *μ*_2_^*^(*t*). The corresponding model M_2_^*^ has the same number of parameters as M. Solving Eq. 2 from Louca & Pennell^2^ with 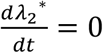 gives:

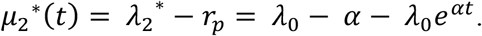

Next, we explore congruent models found by directly fitting *r_p_* to the tree, following the procedure outlined in Louca & Pennel’s SI (section S9), and with the same choice of keeping either *λ* or *μ* constant. We used the *fit_hbd_pdr_on_grid* function from the R package castor^23^ to fit *r_p_*. This function fits *r_p_* assuming a piecewise spline function on a predefined “grid”, i.e. discrete times between which *r_p_* varies as a spline. As we did not find specific recommendations on how to define the grid, we chose evenly-spaced times, and selected their number using the AIC criterion (we note though that the AIC is not a good criterion for rupture detection^25^). We followed the recommendation from the help of the *fit_hbd_pdr_on_grid* function to not use a spline of degree zero (i.e. piecewise constant); we used a spline of degree of 1 (piecewise linear) and 2.

Using a spline of degree 1, we selected 3 times, and the resulting diversification dynamics are shown in Fig. S1. The model that assumes *λ* is constant infers negative extinction rates, and is therefore not plausible. The model that assumes *μ* is constant infers a general tendency for a decline in speciation rates, with a local peak around 7 Myrs ago. This result illustrates again the stability of the general temporal trend (decline) of speciation rates to different fitting approaches. It also illustrates the potential use of congruent models to generate new hypotheses. Here for example, the inferred dynamics suggest to formulate a model with a declining background speciation rate combined with a local burst of speciation around 7 Myrs ago.

Using a spline of degree 2, the AIC criterion selected only 1 time on the grid (i.e a constant *r_p_*), and the resulting diversification dynamics are shown in Fig. S2. The model that assumes *λ* is constant infers negative extinction rates, and is therefore not plausible. The model that assumes *μ* is constant infers a general tendency for a decline in speciation rates that tends to accelerate over time. This result illustrates first that the estimation of r_p_ (and the resulting dynamics of *λ* when assuming *μ* constant) depends on specific hypotheses made when fitting *r_p_* (compare with Fig. S1). Second, it illustrates one more time the stability of the general temporal trend (decline) of speciation rates to different fitting approaches/choices. It also provides another illustration of the potential use of congruent models to generate new hypotheses. Here, the inferred dynamics suggest to formulate a model with a slow decline in speciation rates at the origin of the clade, that accelerates towards the present.

---

**Figure 1.**
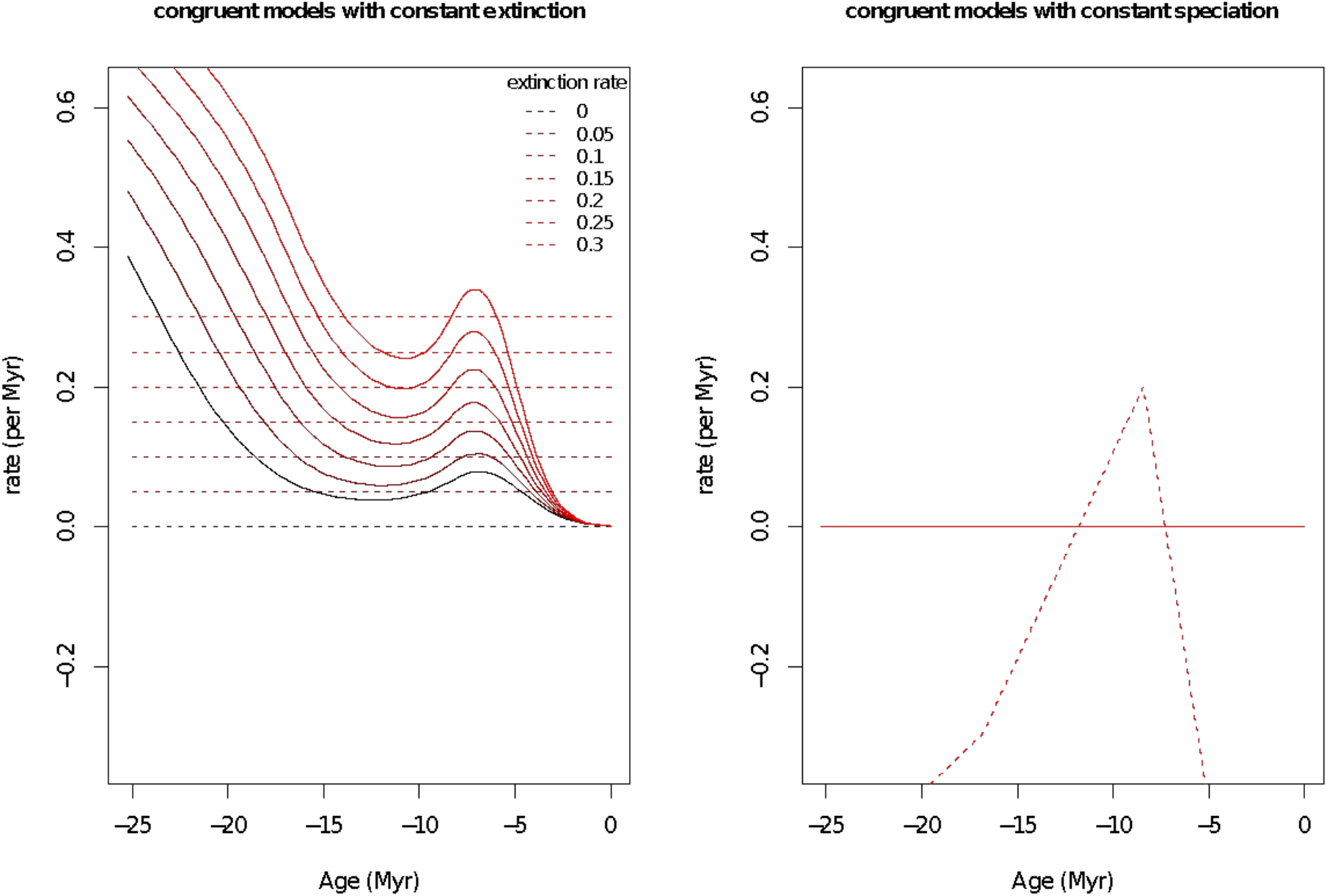
Diversification of the Madagascan vangas as inferred by directly fitting *r_p_*. Here *r_p_* is fitted with a spline of degree 1 (piecewise linear), with 3 evenly-spaced times (selected based on AIC). The curves on the left panel represent the dynamics of speciation (plain line) and extinction (dashed line) obtained by: fixing increasing values of a constant extinction rate (left panel) and fixing the speciation rate to *λ*_0_ (right panel).

**Figure S2.**
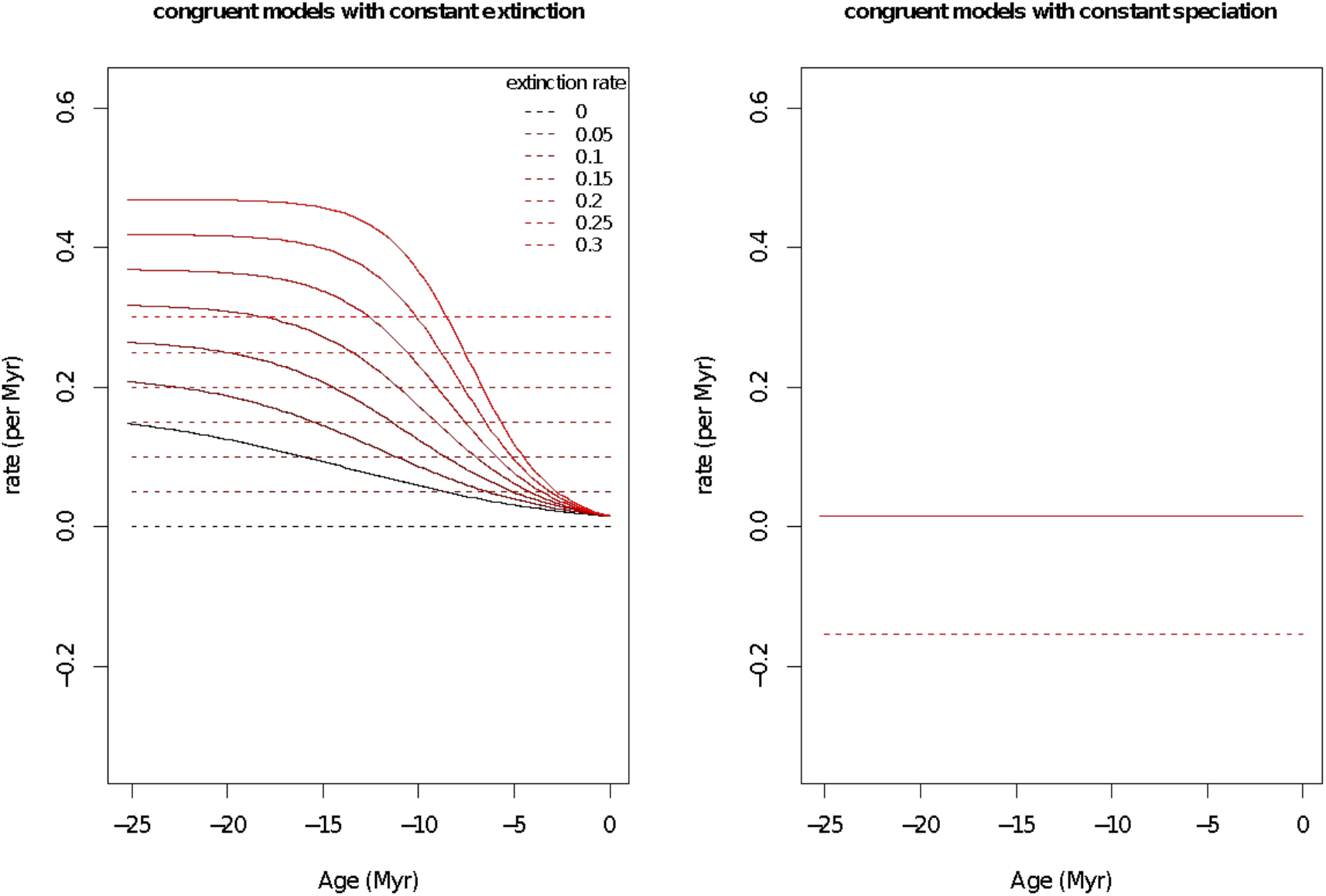
Diversification of the Madagascan vangas as inferred by directly fitting *r_p_*. Here *r_p_* is fitted with a spline of degree 2 with 1 time (selected based on AIC, i.e. *r_p_* is constant). The curves on the left panel represent the dynamics of speciation (plain line) and extinction (dashed line) obtained by: fixing increasing values of a constant extinction rate (left panel) and fixing the speciation rate to *λ*_0_ (right panel).

